# *Grin2a* Ablation Distinctly Reshapes PV+ and SST+ Interneuron circuits in the Medial Prefrontal Cortex to Amplify Gamma Oscillations

**DOI:** 10.1101/2024.12.23.629830

**Authors:** Hassan Hosseini, Sky Evans-Martin, Kevin S. Jones

## Abstract

Schizophrenia (SCZ) is a debilitating mental health disorder marked by cognitive deficits, especially in executive functions, which are often resistant to treatment. The *GRIN2A* gene encodes the GluN2A subunit of N-methyl-D-aspartate (NMDA) receptor. Rare coding mutations in *GRIN2A*, such as protein-truncating variants, increase susceptibility to SCZ by ∼20-fold. However, the effects on network dynamics and the function of GABAergic interneurons (INs) in the medial prefrontal cortex (mPFC) remain unclear. In this study, we use brain slices from *Grin2a* mutant mice to investigate inhibitory synaptic and oscillatory activity driven by parvalbumin-positive (PV+) and somatostatin-positive (SST+) INs. Our findings reveal significant alterations in synaptic currents and inhibitory modulation arise from GluN2A deficiency. Specifically, both heterozygous and knockout *Grin2a* mutants exhibit impaired synaptic transmission and release properties of INs inputs, including changes in release probability and evoked quantal events to layer V pyramidal neurons. Genotype-dependent alterations were observed in the frequency, kinetics, and amplitude of excitatory and inhibitory postsynaptic currents. Immunohistochemical analysis showed an increased density of PV+ and SST+ INs in the *Grin2a* mutants, consistent with an overall increase in inhibitory tone. Additionally, optogenetic stimulation revealed a dynamic shift for inducing gamma-band oscillations through PV+ INs. Overall, our study highlights the essential role of GluN2A-containing NMDA receptors in modulating inhibitory tone and maintaining network stability in the mPFC of adult mice and elucidates a pathophysiological mechanism that might contribute to cognitive deficits often observed in SCZ patients.

## Introduction

Healthy brain function hinges on the precise balance between excitatory and inhibitory (E/I) signaling. Within the medial mPFC, PV+ and SST+ INs represent about 70% of the GABAergic cell population and are essential for tuning network oscillations, integrating synaptic inputs, and supporting cognitive functions such as working memory and decision-making (Kim et al., 2016; Rudy et al., 2011). These executive processes are frequently disrupted in schizophrenia (Arime et al., 2024).

Postmortem analyses of SCZ patient tissue often reveal reduced PV and SST staining (Batiuk et al., 2022; Dienel et al., 2023), and single-nucleus RNA sequencing (snRNAseq) analyses show profound transcriptional changes in SST+ INs (Duncan et al., 2024). Together, these data suggest that SCZ may induce distinct pathophysiological changes in PV+ and SST+ interneuron subpopulations, potentially driving abnormal network dynamics and contributing to the cognitive deficits characteristic of the disorder.

The NMDA receptor (NMDAR) hypofunction model of SCZ has gained traction as a critical framework for understanding the molecular and neural underpinnings of SCZ-related pathophysiology (Coyle et al., 2020; Gao et al., 2022). Loss of function mutations in the *GRIN2A* gene, which encodes the GluN2A subunit of NMDARs, can increase SCZ risk over 20-fold (Harrison & Weinberger, 2005; Moghaddam & Javitt, 2012; Singh et al., 2022). In mice, *Grin2a* hypofunction impairs working memory and synaptic plasticity, and disrupts gamma-band oscillations (GBOs) (Herzog et al., 2023a; Lu et al., 2024; Marquardt et al., 2014; Sakimura et al., 1995) underscoring the pivotal role of GluN2A-containing NMDARs in regulating higher-order cognitive functions and network stability.

Despite extensive research, critical gaps remain regarding how NMDAR hypofunction shapes inhibitory microcircuits at the cellular and synaptic level. Previous studies have established that NMDARs modulate presynaptic GABA release from PV+ INs and postsynaptic plasticity in SST+ INs (Chiu et al., 2018; Pafundo et al., 2018). However, whether *GRIN2A*-associated changes translate into distinct mechanistic alterations within PV+ and SST+ populations is unclear. Notably, while *Grin2a* knockout mice show elevated SST mRNA levels with relatively stable PV mRNA (Lu et al., 2024), the full functional implications of these findings remain elusive. Do PV+ and SST+ INs respond differentially to *Grin2a* ablation, and if so, how do these subtype-specific changes perturb local circuit function and network oscillations?

Addressing these questions is essential for refining SCZ models that often emphasize inhibitory imbalance without dissecting the subtype- and mechanism-specific contributions of PV+ and SST+ INs. Previous work has not fully delineated whether presynaptic and postsynaptic adaptations occur in parallel, nor has it clearly linked these changes to network-level consequences such as altered gamma oscillations. Such insights are crucial, as gamma oscillations are a hallmark of coordinated cortical activity and are frequently disrupted in SCZ (Tanaka-Koshiyama et al., 2020).

In this study, we sought to determine whether *Grin2a* ablation induces distinct, mechanistically distinct changes in PV+ and SST+ IN function within the adult mPFC and to understand how these alterations influence network oscillations. We combined optogenetics, whole-cell patch-clamp electrophysiology, multi-electrode array (MEA) recordings, and immunohistochemistry to dissect presynaptic versus postsynaptic changes in inhibitory circuits and their impact on GBOs. By doing so, we establish that *Grin2a* hypofunction diverges along interneuron subtype lines: PV+ IN alterations predominantly arise from disrupted presynaptic calcium dynamics and asynchronous GABA release, while SST+ IN changes manifest primarily at the postsynaptic level. Crucially, these distinct mechanisms converge to modify E/I balance and amplify gamma oscillations in the mPFC.

Our findings provide a nuanced mechanistic framework that links a well-characterized genetic risk factor for SCZ—*Grin2a* dysfunction—to specific synaptic and oscillatory disruptions. This refined perspective not only helps reconcile discrepancies with previous studies (e.g., developmental differences in circuit maturation or methodological variations) but also advances the field by highlighting the importance of timing, location, and cellular specificity in shaping SCZ-related phenotypes. Ultimately, these insights may guide targeted therapeutic strategies to restore inhibitory balance and network stability, offering more precise interventions to mitigate cognitive impairments observed in SCZ.

## Methods

### Animals

All animal procedures were approved by the University of Michigan’s Institutional Animal Care and Use Committee (IACUC) and conformed to NIH Guidelines for animal use. *Grin2a*^KO^ mice (Riken B6;129S-Grin2a<tm1Nak>; RBRC02256) were intercrossed or backcrossed to C57BL/6J mice to generate *Grin2a*^HET^ or *Grin2a*^KO^ offspring (Kadotani et al., 1996). Genotypes were confirmed by real-time PCR (Transnetyx, Cordova, TN). Male mice were exclusively used in this study to eliminate the variability introduced by the female estrous cycle, thereby ensuring more consistent and reliable results.

### Stereotactic Surgeries

Male mice (7–8 weeks old) were anesthetized with 4–5% isoflurane in an induction chamber, then transferred to a stereotactic frame (Kopf Instruments, model 1400) and maintained at 1–2% isoflurane via a nose cone. The head was stabilized, the scalp shaved and sterilized, and a 1 cm incision made along the anterior-posterior axis. Bilateral craniotomies were performed above the mPFC (coordinates: AP: +1.9 mm, ML: ±0.3 mm, DV: −1.8 mm). AAV1-SST-hChR2(H134R)-EYFP (Karl Deisseroth lab, Nadia Andini (Mattis et al., 2014)) or pAAV1-S5E2-ChR2-mCherry (AddGene) was injected bilaterally (110 nl) using a Nanoject II (Drummond Scientific, USA). The virus was allowed to diffuse into the tissue for an additional 5-7 min after the end of injection before the pipette withdrawal. The incision was then sealed with surgical glue. Mice received carprofen (5 mg/kg, s.c.) immediately after surgery and again the following day. Mice were monitored for 7-10 days post-treatment.

### Acute Slice Preparation

Coronal mPFC slices (300 μm thick) were prepared from 12- to 15-week-old male mice using established protocols (Hosseini et al., 2024). Briefly, brains were rapidly extracted after isoflurane anesthesia and decapitation, then immersed in ice-cold, carbogenated slicing solution. Slices were cut using a vibratome and incubated in recovery solution prior to recordings.

### MEA Slice Electrophysiology

Local field potentials (LFPs) were recorded from mPFC coronal slices following previously described methods (Hosseini et al., 2024). Recordings were conducted after a 15-minute baseline period. Optical stimulation (470 nm, 10 ms pulse width) was delivered using a digital micromirror device (Polygon-400, Mightex) through a 10x objective (Olympus) at frequencies ranging from 1 Hz to 128 Hz to selectively activate PV+ or SST+ INs. Optical power output was maintained at 1 mW/mm².

### Whole-cell Patch Clamp Electrophysiology

Whole-cell patch-clamp recordings were obtained from visually identified putative layer V pyramidal neurons based on their location, size, and morphology in the pre-limbic (PrL) region of the mPFC. Borosilicate glass pipettes (3–5 MΩ) were filled with either a K⁺- or Cs⁺-based internal solution, as appropriate for recording excitatory or inhibitory postsynaptic currents, respectively. Signals were amplified (MultiClamp 200B), filtered at 8 kHz, and digitized at 20 kHz (Digidata 1550A, Molecular Devices).

Spontaneous EPSCs (sEPSCs) and IPSCs (sIPSCs) were recorded in standard ACSF at −70 mV and 0 mV, respectively. Miniature EPSCs (mEPSCs) and IPSCs (mIPSCs) were recorded under similar conditions with the addition of 1 µM TTX. To evaluate synaptic depression and asynchronous IPSC (asIPSC) activity, 40 Hz trains of optical stimuli (1 s) were delivered to PV+ INs. Frequency response curves were generated by normalizing the asIPSC area (2 s post-stimulation) to the mean area of 3 pre-tetanic IPSCs. EGTA-AM (100 µM) was used to chelate residual Ca²⁺ and isolate asIPSC activity (Hefft & Jonas, 2005; Jensen et al., 2000).

### Optogenetic Stimulation

Optogenetic activation of PV+ and SST+ INs in the PrL region was achieved using a 470 nm LED (0.5–1 mW/mm²; Thorlabs, Newton, NJ) controlled by a digitally-controlled driver. The light was delivered through a 40x objective (Olympus BX51W). Stimuli were adjusted to synchronize responses and eliminate polysynaptic activity. A 1-ms pulse duration was used, and series resistance was compensated by 70% following patch breakthrough. To measure paired-pulse ratios (PPRs) the amplitude of the second evoked IPSC (eIPSC2) was divided by the first (eIPSC1). For short inter-stimulus intervals (ISI), the prolonged decay of eIPSC1 was subtracted from eIPSC2 to obtain accurate measurements. Evoked miniature IPSCs (mIPSCs) were recorded by optogenetically stimulating axons proximal to recorded pyramidal neurons in the presence of TTX, APV, CNQX, and 4-AP adapted from Miska et al. (2018). Neurons were voltage-clamped at 0 mV using an improved Cs⁺-based internal solution. After baseline acquisition, neurons were stimulated with continuous 470 nm light for 5 s. Stimulus power was adjusted to ensure evoked frequencies were significantly above spontaneous activity, with clearly discernible individual events. The following criteria were used to determine data inclusion: Series resistance below 17 MΩ at baseline, with fluctuations <15% during recording. Holding leak current < −200 pA, with deviations <50 pA. Post hoc immunohistochemistry confirmed the recorded neurons’ locations relative to the viral injection site, using biocytin staining to verify their morphology and connectivity.

### Histology

Tissue preparation, permeabilization, and antibody incubation were performed as described previously (Hosseini et al., 2024). Primary antibodies: Guinea pig anti-SST (SYSY, #366004, 1:1000), rabbit anti-PV (Swant, #PV27a, 1:1000), mouse anti-GAD67 (Millipore, #MAB5406, 1:1000). Secondary antibodies: Alexa-594 donkey anti-rabbit (Invitrogen #A21207), Alexa-647 goat anti–guinea pig (Life Technologies #A21450), Alexa-488 donkey anti-mouse (Life Technologies #A21202). Images were analyzed using ImageJ (https://imagej.nih.gov/ij/), counting cells in 3–4 sections (60 μm) rostrocaudally per mouse.

### MEA Data Processing and Statistical Analysis

The data was processed using Python. A notch filter removed 60 Hz noise. The 30–100 Hz range was designated as gamma (Colgin et al., 2009). Absolute power within each frequency band was calculated by integrating the power spectrum. Data was processed with custom Python scripts. LFPs were low-pass filtered at 500 Hz and downsampled to 1 kHz. Multi-taper spectral analysis was used for power estimation. Relative power changes were computed as ((induced − baseline)/baseline). The 1 Hz optogenetic stimulation response amplitude was measured as peak LFP amplitude. All analysis code is available at https://github.com/NeuroDataa/MEA_Analysis.

### Patch Clamp Data Processing and Statistical Analysis

Data was analyzed in Python. Passive and active properties were extracted using the eFEL package (Ranjan et al., 2024), with additional custom Python scripts available at https://github.com/NeuroData/PatchClampAnalysis. For sEPSC and sIPSC detection, traces were filtered at 1 kHz, and events were identified using template matching in pClamp (Clampfit). Templates were defined from averages of >100 events.

## Results

### The Density of PV+ and SST+ INs is Increased in the PrL of *Grin2a* Mutants

Previous reports have shown that *Grin2a* mutation increases cortical GABAergic interneuron (IN) density (Camp et al., 2023; Lu et al., 2024). To determine whether this effect targets specific IN subtypes, we stained mPFC sections from WT, *Grin2a*^HET^, and Grin2a^KO^ mice for PV+, SST+, and GAD67+ markers (**Fig. 1A-C**). Both *Grin2a*^HET^ and Grin2a^KO^ mice exhibited significantly higher PV+ and SST+ IN densities in PrL layer II/III and V compared to WT (**Fig. 1D-E**). However, their proportions relative to total GAD67+ cells were unchanged (**Fig. 1F, G).** This suggests that *Grin2a* ablation broadly increases GABAergic IN populations rather than a particular subtype.

**Figure 1.**
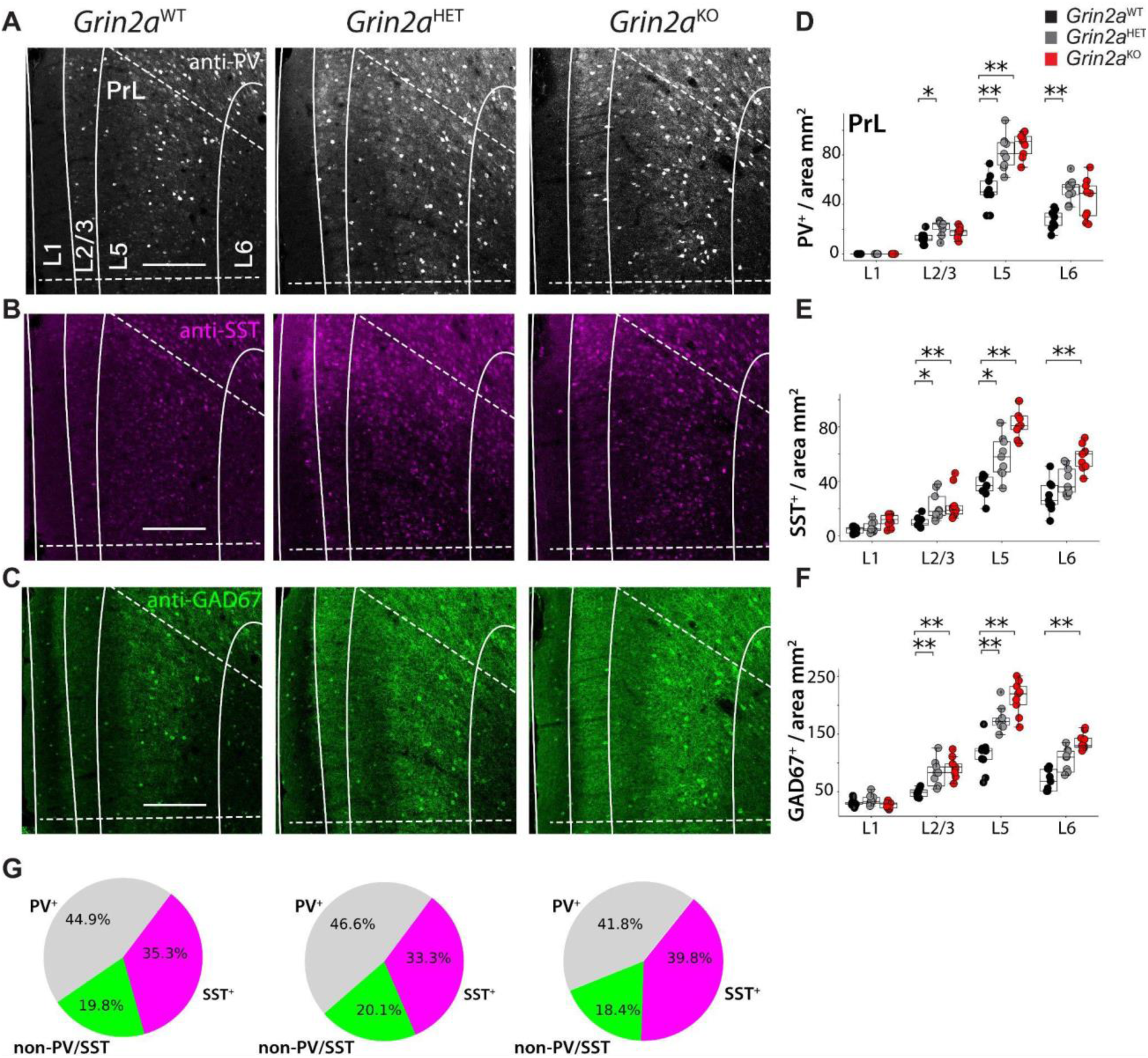
GABAergic IN Density in the PrL Region of *Grin2a* Deficient Mice. (**A–C**) Representative confocal micrographs of PrL sections from WT, *Grin2a*^HET^, and *Grin2a*^KO^ mice stained for PV+ (A), SST+ (B), and GAD67+ (C). (**D–F**) Quantification of PV+, SST+, and GAD67+ cell densities across cortical layers (L1–L6) in WT and *Grin2a* mutants. *Grin2a*^HET^ and *Grin2a*^KO^ mice show significantly increased PV+ and SST+ cell densities, particularly in L2/3 and L5, compared to WT. GAD67+ cell density also increases, indicating a broad rise in GABAergic interneurons rather than a subtype-specific expansion. (**G**) Pie charts illustrating the ratio of PV+ and SST+ cells to total GAD67+ cells in PrL L5. Despite increased absolute densities, the relative proportions of PV+ and SST+ cells to total GAD67+ INs remain stable. Data are presented as mean ± SEM from 3 mice per genotype, with 3–4 sections (60 μm) per mouse. Statistical significance was determined using the Kruskal-Wallis test: *p < 0.05, **p < 0.01. Scale bars, 200 μm.

### The E/I Balance in PrL layer V Pyramidal Neurons is Reduced in *Grin2a* Mutants

To assess whether increased GABAergic IN density influences excitatory-inhibitory (E/I) balance in the mPFC, we recorded whole-cell currents from layer V pyramidal neurons in the PrL (**Fig. 2A**). While *Grin2a*^HET^ and *Grin2a*^KO^ neurons showed sEPSC frequencies similar to WT (**Fig. 2B**), their mean sEPSC amplitudes were reduced by ∼45% (**Fig. 2C)**. In contrast, sIPSC frequency was doubled in *Grin2a* mutants (**Fig. 2B**), with no change in sIPSC amplitude or decay kinetics (**Fig. 2D**). We analyzed the distribution of sIPSC amplitudes. We observed a rightward shift in the median for both *Grin2a*^HET^ and *Grin2a*^KO^ neurons, reflecting a higher frequency of large-amplitude sIPSCs (**Fig. 2E**). Additionally, mIPSC frequency was significantly elevated in *Grin2a* mutant neurons compared to WT, whereas mEPSC frequency remained unchanged (**Fig S1A-C**). Despite pronounced alterations in synaptic inputs, the intrinsic properties of pyramidal neurons in *Grin2a* mPFC were largely preserved. The only notable exceptions were increased spike amplitude and width (**Fig. S2A-E**). These results demonstrate that *Grin2a* ablation significantly disrupts the E/I balance in PrL layer V pyramidal neurons by reducing excitatory drive and increasing inhibitory input.

**Figure 2.**
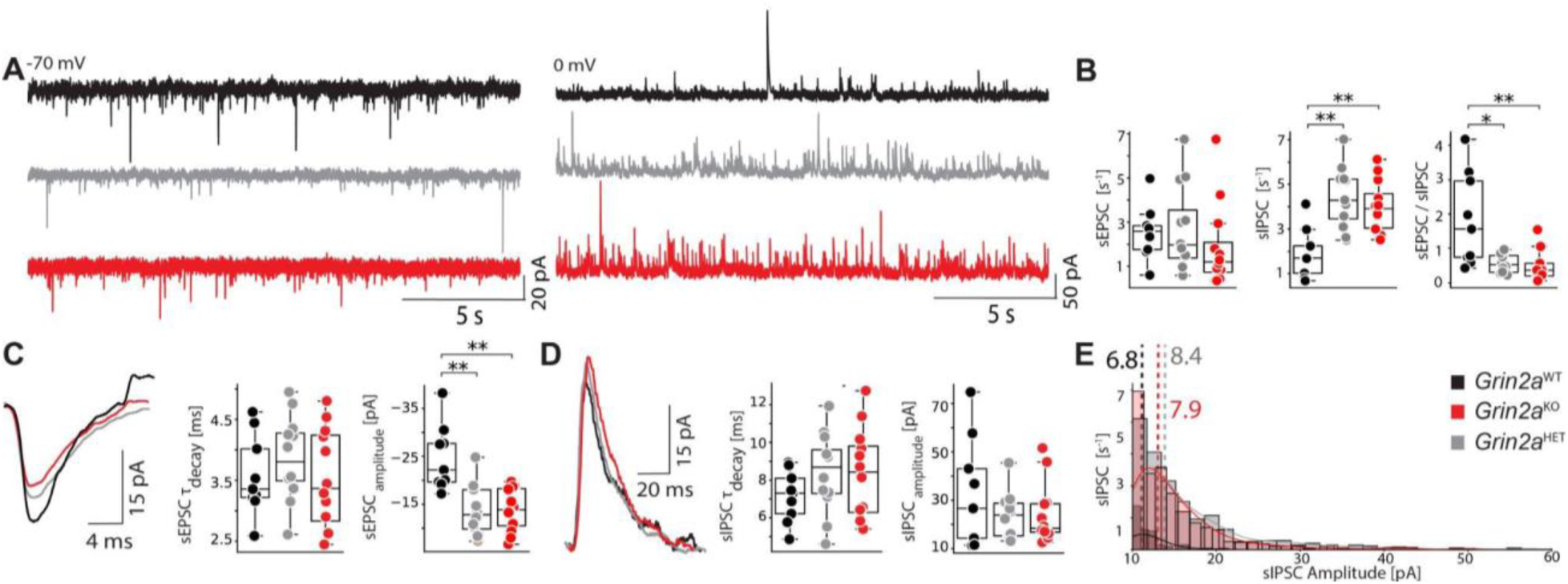
Spontaneous Excitatory and Inhibitory Currents in *Grin2a* Deficient Mice. (**A**) Representative electrophysiological traces of spontaneous excitatory postsynaptic currents (sEPSCs) at −70 mV and spontaneous inhibitory postsynaptic currents (sIPSCs) at 0 mV in layer V pyramidal neurons from WT, *Grin2a*^HET^, and *Grin2a*^KO^ mice. (**B**) Box plots showing mean sEPSC frequency (Hz) and mean sIPSC frequency (Hz) across genotypes. The ratio of sEPSC to sIPSC frequencies is significantly decreased in *Grin2a* mutants. (**C**) Representative traces and quantification of sEPSC decay tau (ms) and amplitude (pA). (**D**) Representative traces and quantification of sIPSC decay tau (ms) and amplitude (pA). (**E**) Histogram of sIPSC frequency versus amplitude for each genotype, with median amplitude indicated by vertical lines. Data represent mean ± SEM from 10–15 cells per genotype. Statistical significance was determined using the Kruskal-Wallis test: *p < 0.05, **p < 0.01, ***p < 0.001.

### Presynaptic Transmission from PV+ INs is Altered in *Grin2a* Mutants

Building on previous reports that *Grin2a* ablation alters electrical properties of PV+ INs (Camp et al., 2023), we examined the impact on PV+ synaptic output. Using optogenetic stimulation, we elicited PV-driven IPSCs in layer V of PrL pyramidal neurons (**Fig. 3A**). Although the amplitude of evoked IPSCs (eIPSCs) did not differ significantly between genotypes, decay times were prolonged 2.3-fold in *Grin2a*^HET^ neurons and 2.6-fold in *Grin2a*^KO^ neurons compared to WT (p = 0.002 and p = 0.01, respectively) (**Fig. 3B, C**). To determine if prolonged decay arose from alterations in presynaptic function, we measured the PPR of PV-driven IPSCs. PPR was impaired in *Grin2a* mutants, with the most pronounced differences observed at a 100 ms interstimulus interval (**Fig. 3D, E**). This data suggests that *Grin2a* ablation increases PV-driven presynaptic release probability, indicating a disruption in Ca²⁺-dependent release efficacy.

**Figure 3.**
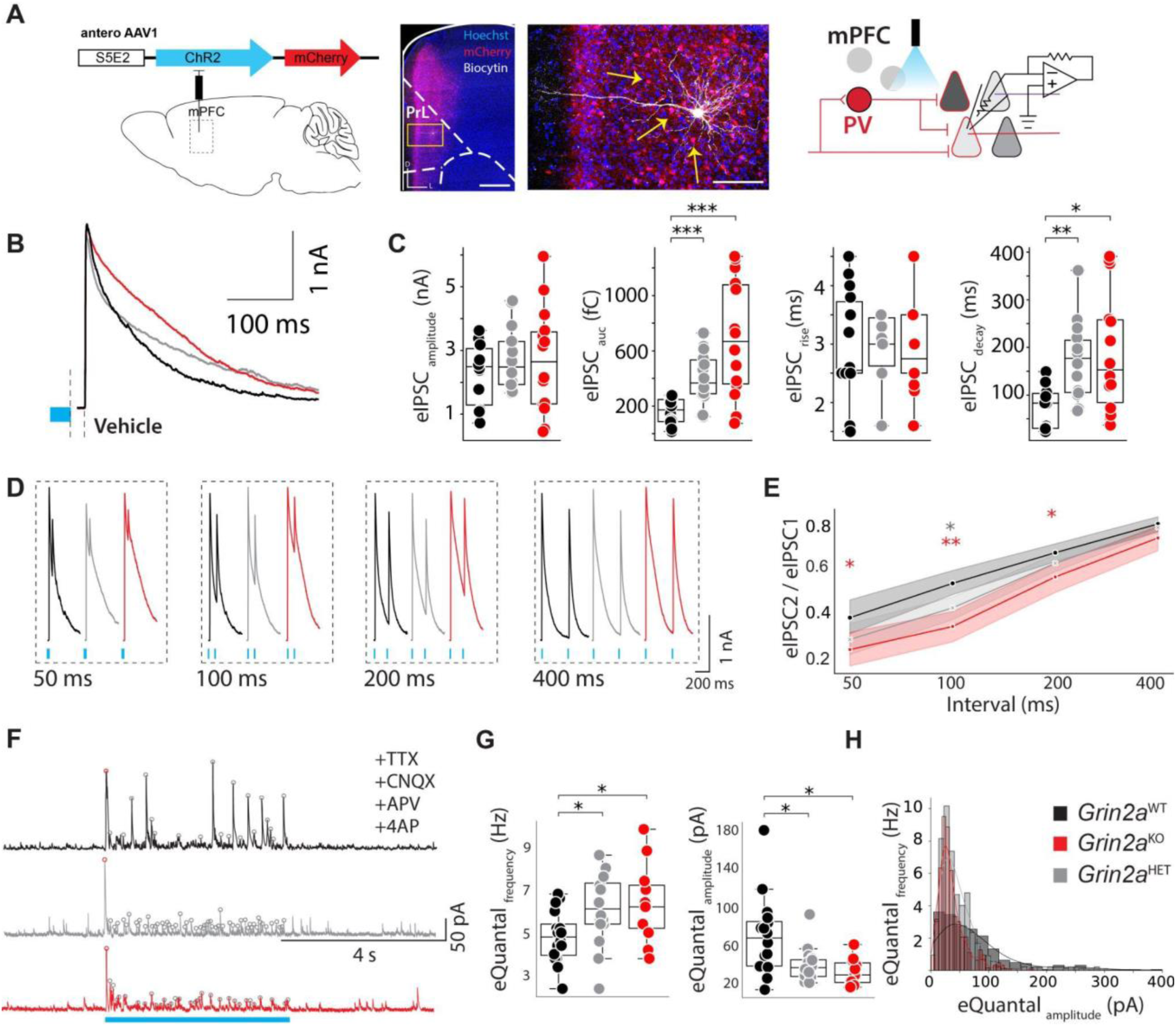
Synaptic and Release Properties of Evoked IPSCs from PV+ INs Input to Layer V Pyramidal Neurons in PrL. (**A**) Schematic of viral injection (pAAV-S5E2-ChR2-mCherry) into mPFC (left), confocal micrograph of a biocytin-filled recorded layer V pyramidal cell (middle), and schematic of whole-cell patch-clamp recording from a pyramidal cell (right). Scale bars, 500 μm, and 100 μm, respectively. (**B**) Representative traces of PV-driven evoked IPSCs (eIPSCs) in *Grin2a* mutants. (**C**) Summary statistics of eIPSC amplitude (nA), area under the curve (pA·ms), rise tau (ms), and decay tau (ms) across genotypes. (**D**) Representative traces of paired-pulse ratio (PPR) for eIPSCs. (**E**) Summary statistics of PPR at different interstimulus intervals (ms) across genotypes. (**F**) Representative traces of PV-evoked quantal events during 5 s stimulation. (**G**) Evoked quantal frequency (Hz) and amplitude (pA) are quantified across genotypes. (**H**) Scatter plot of evoked quantal frequency versus amplitude for each genotype. Data represent mean ± SEM from 10–15 cells per genotype. Statistical significance was determined using the Kruskal-Wallis test:*p < 0.05, **p < 0.01, ***p < 0.001

### Presynaptic Mechanisms Underlying Altered PV+ Transmission

To distinguish action potential (AP)-dependent from AP-independent changes, we measured PV-driven quantal release under conditions that isolate miniature events (Miska et al., 2018). In *Grin2a* mutant slices, quantal amplitude decreased by ∼48%, while quantal frequency increased by ∼30% relative to WT (**Fig. 3F-H)**. These shifts suggest that *Grin2a* ablation alters presynaptic function, affecting both the magnitude and probability of release events.

### Asynchronous GABA Release from PV+ INs is Enhanced in *Grin2a* Mutants

The slow IPSC decay observed in Grin2a PV+ INs may stem from elevated presynaptic calcium accumulation (Chamberland et al., 2020). To test this, we applied EGTA-AM, a membrane-permeable calcium chelator. EGTA-AM had minimal effect on WT slices but significantly accelerated IPSC decay in *Grin2a*^HET^ and *Grin2a*^KO^ slices, restoring kinetics to WT levels (**Fig. 4A, B**). Because asynchronous GABA release often arises from residual presynaptic calcium, we next examined asIPSCs following 40 Hz tetanic stimulation, a paradigm designed to maximize asynchronous release (**Fig. S3A, B**). AsIPSCs were substantially larger in *Grin2a* mutants, and EGTA-AM reduced these events by roughly twice as much in mutants as WT (**Fig. 4C-D)**. Moreover, *Grin2a* ablation slowed the onset of synaptic depression during tetanic stimulation, and EGTA-AM countered this effect more strongly in mutants than WT neurons (**Fig. S3C, D**). These findings align with models in which elevated presynaptic calcium prolongs asynchronous release and delays synaptic depression (Dittman et al., 2000).

**Figure 4.**
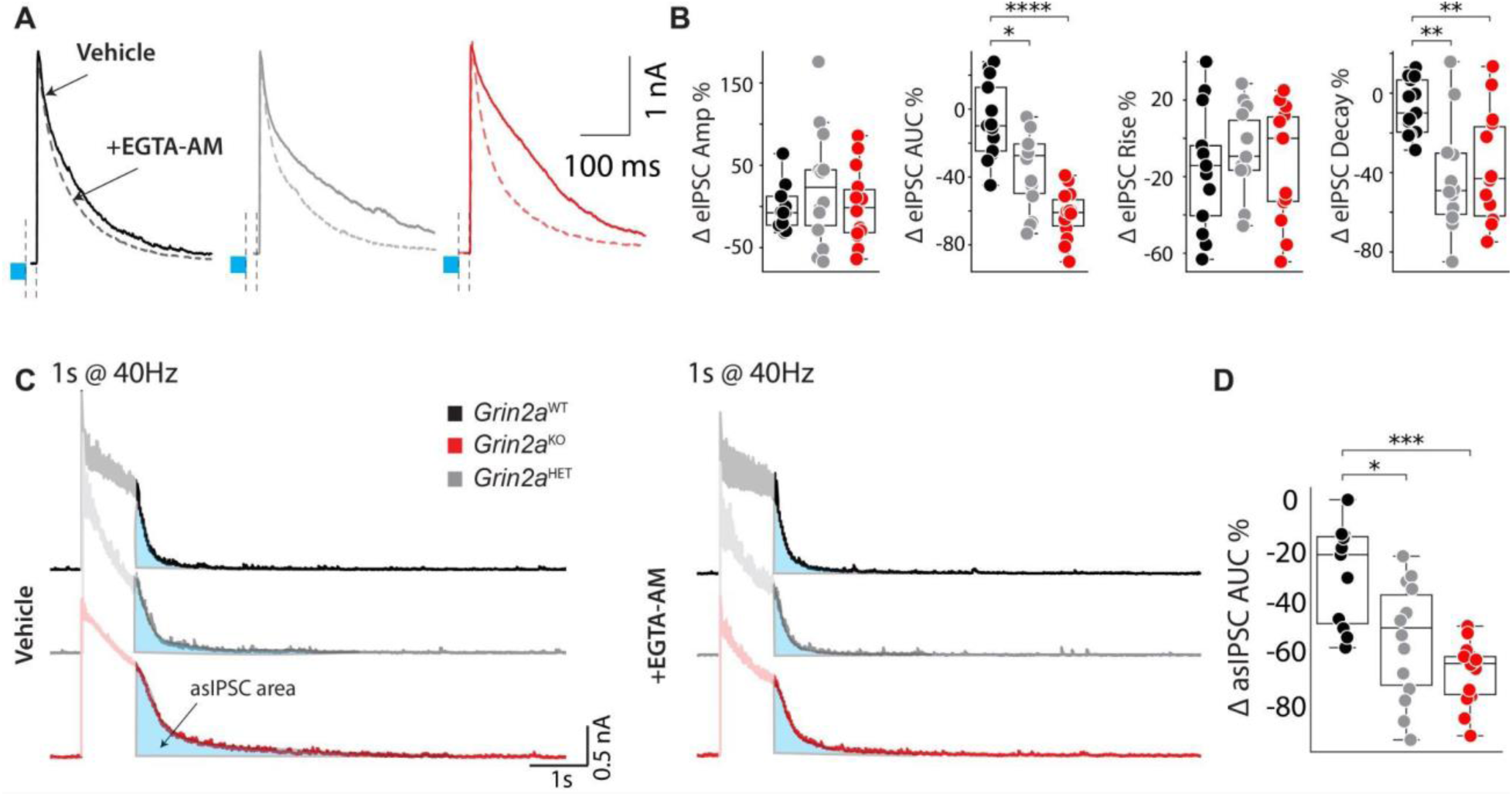
PV+ Opto-Evoked Post-Tetanic asIPSC in PrL Layer V Pyramidal Neurons in *Grin2a* Mutants. (**A**) Representative averaged trace of PV-driven eIPSC amplitudes under baseline conditions and in the presence of EGTA-AM (100 μM). (**B**) Summary statistics of percent change in eIPSC peak amplitude (%), area under the curve (AUC; pA·ms), rise tau (ms), and decay tau (ms) across genotypes. (**C**) Representative averaged trace of asynchronous IPSCs (asIPSCs) during 40 Hz tetanic stimulation for 1 s. (**D**) Summary statistics of fold change percentage in asIPSCs after 40 Hz train stimulations (%). Data represent mean ± SEM from 10–15 cells per genotype. Statistical significance was determined using the Kruskal-Wallis test: *p*t:*p < 0.05, **p < 0.01, ***p < 0.001.

### PV-Driven iGBOs are Enhanced in *Grin2a* Mutants

Fast-spiking, PV+ INs are critical drivers of GBOs (Cardin et al., 2009; Zarnadze et al., 2016). To examine how *Grin2a* ablation affects the inhibitory regulation of cortical circuits, we recorded LFPs from mPFC slices while optogenetically stimulating PV+ INs with increasing frequency (**Fig. 5A**). There was no difference in the amplitudes of PV+ IN-evoked LFPs of WT or *Grin2a* mutants (**Fig. 5B, D**). Stimulus frequencies of ≥32 Hz increased the power of GBOs in layer II/III or layer V of WT and both *Grin2a* mutants (**Fig. 5C, E**). High-frequency stimulation increased the power of iGBOs in *Grin2a*^KO^ slices by 2.5-fold compared to WT slices in layer II/III (p = 0.019 at 64 Hz and p = 0.015 at 128 Hz; (**Fig. 5C**) and it decayed ∼2-fold slower than WT following maximum stimulation (**Fig. S5A-B**). This is consistent with elevated residual calcium in PV+ terminals prolonging inhibitory signaling. These findings suggest that *Grin2a* deficiency amplifies PV-driven iGBOs, indicating altered inhibitory modulation of network oscillations.

**Figure 5.**
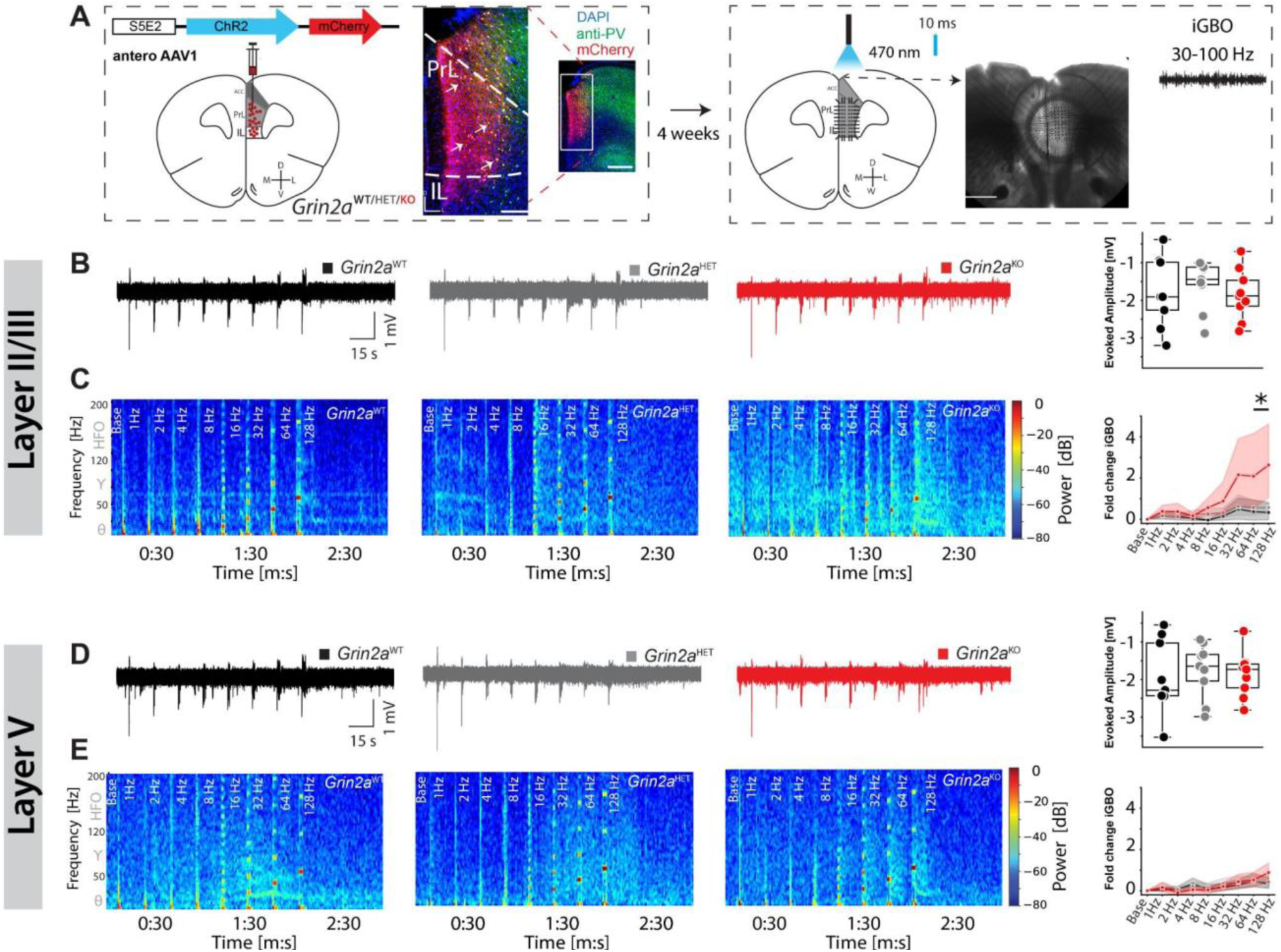
PV-Driven iGBOs in mPFC Sections from *Grin2a* Mutant Mice. (**A**) Schematic of viral injection (pAAV-S5E2-ChR2-mCherry) into mPFC (left) and placement on a perforated MEA, along with a bright-field micrograph of an *ex vivo* mPFC slice. Scale bars, 300 μm and 500 μm. (**B**) Example LFP recordings from layer II/III mPFC following opto-stimulation of PV+ INs, with boxplots of evoked amplitudes (mV) at 1 Hz across genotypes. (**C**) Representative layer II/III spectrograms following opto-stimulation of PV+ INs and line graphs of iGBO power over 5 seconds post-stimulation at various frequencies. Fold change in iGBO power is shown for each frequency across genotypes. (**D**) Example LFP recordings from layer V mPFC following opto-stimulation of PV+ INs, with boxplots of evoked amplitudes (mV) at 1 Hz across genotypes. (**E**) Representative spectrograms of layer V following opto-stimulation of PV+ INs and line graphs of iGBO power 1 second post-stimulation at various frequencies. Fold change in iGBO power is shown for each frequency across genotypes. Data represent mean ± SEM from 8–9 mPFC slices from 3 mice per genotype. Statistical significance was determined using the Kruskal-Wallis test with Bonferroni correction for multiple comparisons::*p < 0.05, **p < 0.01, ***p < 0.001

### SST-Driven IPSCs are Enhanced in *Grin2a* Mutants

NMDAR activation has been implicated in the postsynaptic regulation of SST-driven inhibition (Chiu et al., 2018). To assess how *Grin2a* ablation affects SST-mediated synaptic transmission in the PrL mPFC, we measured SST-driven eIPSCs in layer V pyramidal neurons (**Fig. 6A**). The amplitude of SST-driven eIPSCs were significantly larger in *Grin2a*^HET^ and *Grin2a*^KO^ mice compared to WT, and the eIPSC decay kinetics of *Grin2a*^KO^ slices were slower (**Fig. 6B, C)**. In contrast, short-term plasticity and Ca²⁺-dependent release properties at SST synapses remained unchanged **(Fig. 6D-E; Fig. S4A-B)**. These results suggest that *Grin2a* deficiency alters SST synapse function primarily via postsynaptic mechanisms, increasing inhibitory drive without disturbing release dynamics.

**Figure 6.**
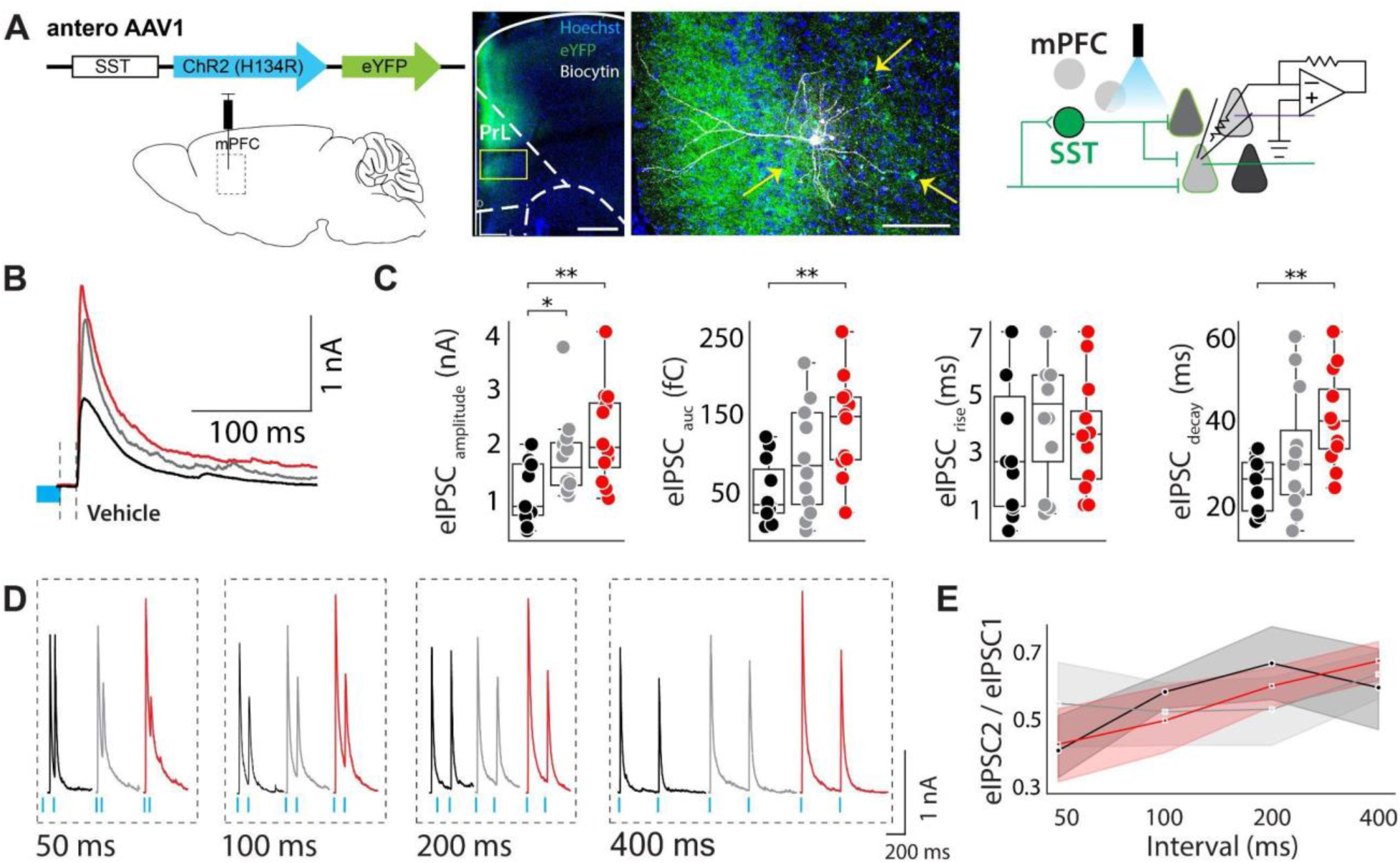
Synaptic and Release Properties of Evoked IPSCs from SST+ IN Input to Layer V Pyramidal Neurons in PrL. (**A**) Schematic of viral injection (pAAV-SST-ChR2-eYFP) into mPFC (left), confocal images of a biocytin-filled recorded pyramidal cell (middle), and schematic of whole-cell patch-clamp recording from an L5 pyramidal cell (right). Scale bars, 500 μm and 100 μm. (**B**) Representative averaged trace of SST-driven eIPSCs under baseline conditions. (**C**) Summary statistics of baseline eIPSC peak amplitude (nA), area under the curve (AUC; pA·ms), rise tau (ms), and decay tau (ms) across genotypes. (**D**) Representative traces of eIPSC PPR across genotypes. (**E**) Line graphs of PPR at different interstimulus intervals (ms) across genotypes. Data represent mean ± SEM from 10–15 cells per genotype. Statistical significance was determined using the Kruskal-Wallis test:s::*p < 0.05, **p < 0.01, ***p < 0.001

### SST-driven iGBOs are Unchanged in *Grin2a* Mutants

Optogenetic stimulation of ChR2-expressing SST+ INs induces iGBOs in hippocampal slices (Antonoudiou et al., 2020). Here, we stimulated ChR2-expressing SST+ INs and recorded LFPs from the PrL of mPFC slices from *Grin2a* mutants (**Fig. 7A**). In layer II/III, the amplitude of LFPs evoked in *Grin2a^HET^* slices was ∼70% greater than in WT (**Fig. 7B**). However, despite the increased LFP amplitude, the iGBO threshold (∼32 Hz) and power-frequency relationship remained unchanged across genotypes (**Fig. 7C)**. Similarly, in layer V, *Grin2a*^HET^ slices exhibited LFP amplitudes, but iGBO power profiles were comparable to WT (**Fig. 7D, E**). Thus, while *Grin2a* ablation enhances SST-driven synaptic input, it does not disrupt iGBO frequency or power. The preserved short-term plasticity and Ca²⁺-dependent release at SST synapses likely maintains synchronous GABA release, stabilizing iGBO frequency and power.

**Figure 7.**
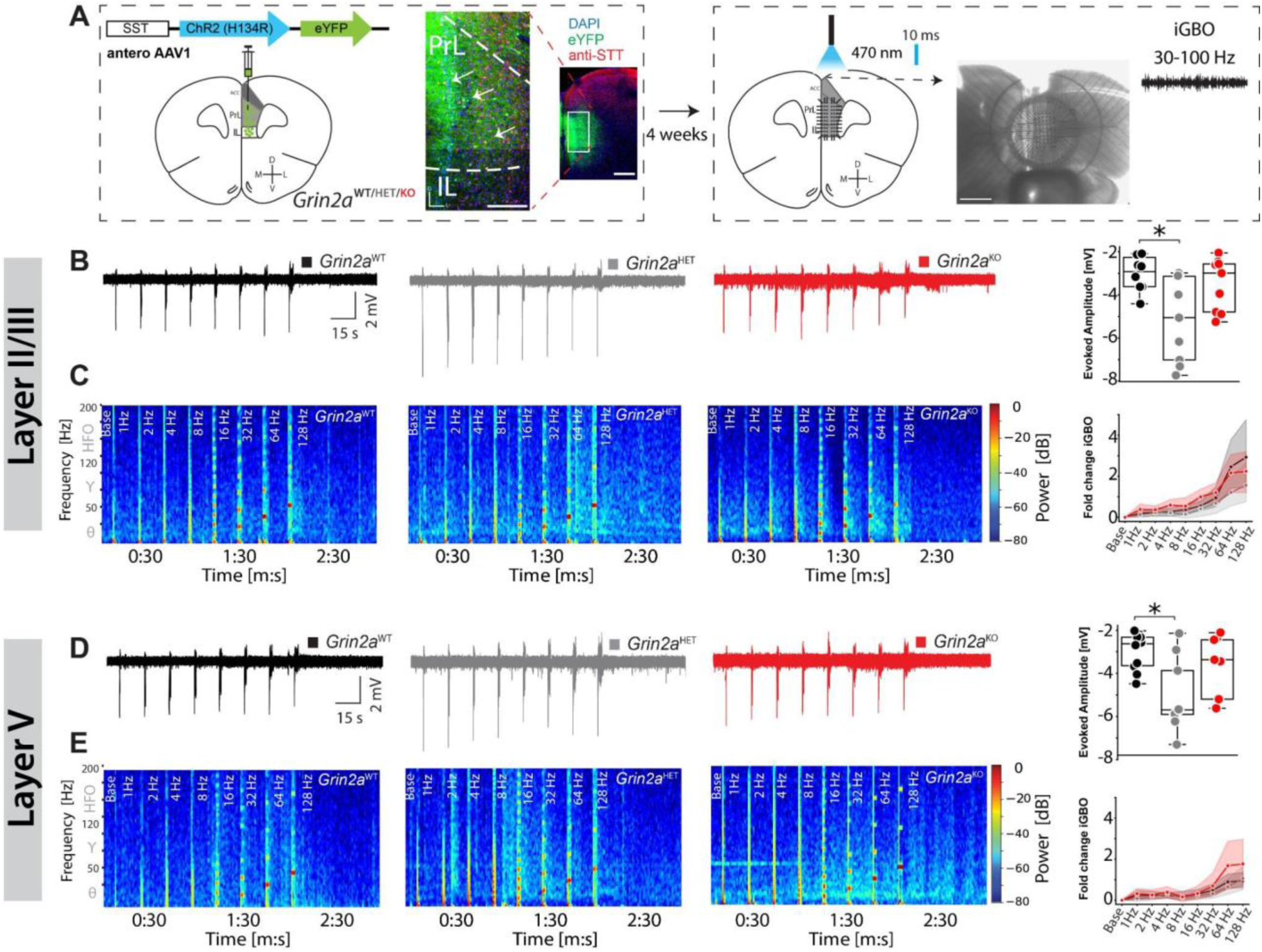
SST-Driven Inhibitory Oscillations in *Grin2a* Mutants Maintain Frequency Relationships. **(A)** Experimental setup for optogenetic stimulation of SST+ interneurons in the mPFC. Left: Schematic of AAV-ChR2-mCherry injection targeting SST+ cells and representative confocal images illustrating viral expression in the PrL region (scale bars: 300 μm and 500 μm). Right: A schematic of the ex vivo perforated multi-electrode array (pMEA) recording configuration and an example of optogenetic illumination (470 nm, 10 ms pulses) used to evoke inhibitory gamma-band oscillations (iGBOs). **(B–C)** LFP recordings (B) and corresponding spectrograms (C) from layer II/III show that optogenetic stimulation of SST+ INs elicits robust inhibitory responses in all genotypes. Although *Grin2a*^HET^ and *Grin2a*^KO^ slices display similar or enhanced initial response amplitudes compared to WT, the frequency-dependent changes in iGBO power remain relatively modest. Over a range of stimulus frequencies, iGBO power increases in all genotypes but does not exhibit the pronounced frequency-dependent elevation observed with PV+ stimulation (see Fig. 5). **(D–E)** Recordings (D) and spectrograms (E) from layer V similarly reveal SST-driven iGBOs that are slightly enhanced in *Grin2a* mutants compared to WT, but without the dramatic, frequency-specific amplification noted in PV+ networks. While SST+ stimulation increases inhibitory input and can alter amplitude, it does not substantially reshape the frequency response curve of gamma oscillations as observed with PV+ circuits. Data represent mean ± SEM from 8–9 mPFC slices from 3 mice per genotype. Statistical significance was determined using the Kruskal-Wallis test with Bonferroni correction for multiple comparisons, with p < 0.05 (*).

## Discussion

This study highlights that GluN2A-containing NMDARs are critical for maintaining interneuron function and stable network dynamics in the mPFC. *Grin2a* ablation increases PV+ and SST+ interneurons (INs) density, tipping the E/I balance toward inhibition and reducing excitatory drive onto layer V pyramidal neurons. In turn, these changes amplify gamma-band oscillations, especially in upper cortical layers. Together, these findings show how GluN2A dysfunction reshapes inhibitory tone, informing our understanding of NMDAR hypofunction in neuropsychiatric conditions like schizophrenia.

### Differential Impact on PV+ and SST+ INs

*Grin2a* ablation affects PV+ and SST+ INs differently. PV+ cells show prolonged IPSC decay and more asynchronous GABA release, consistent with presynaptic calcium dysregulation. Elevated paired-pulse ratios and quantal release frequency indicate a higher vesicle fusion probability. By contrast, SST+ INs primarily undergo postsynaptic changes, evidenced by increased IPSC amplitudes without shifts in short-term plasticity or calcium-dependent release.

### Impact on Cortical Oscillations

Presynaptic calcium dysregulation in PV+ INs appears central to the enhanced gamma oscillations in layer II/III. Prolonged paired-pulse depression and increased asynchronous GABA release disrupt synaptic timing, reducing the precision of inhibitory control. Consequently, elevated gamma power may represent a maladaptive synchronization that contributes to cognitive deficits in SCZ.

### Alignment with Prior Studies

These results align with prior work linking *GRIN2A* mutations to neurodevelopmental and cognitive deficits (Singh et al., 2022). Consistent with Kinney et al. (2006) and Hosseini et al. (2024), who identified impaired PV+ activity and altered E/I balance in the hippocampus, we now show similar disruptions in the mPFC. Herzog et al. (2023) further demonstrated elevated cortical GBO power in *Grin2a*^HET^ mice, paralleling SCZ-related electrophysiological changes. Such concordance reinforces the translational value of these models. Moreover, recent genetic and transcriptomic data (Duncan et al., 2024) highlighting SST+ IN involvement in SCZ underscores the subtype-specific contributions to network dysfunction.

A subset of our findings contrasts with Lu et al., who reported no changes in sIPSC frequency or amplitude and increased sEPSC frequency in *Grin2a* mutants (Lu et al., 2024). These discrepancies may arise from differences in developmental stages between studies as inhibitory networks in the PFC mature gradually into adulthood (Caballero et al., 2014). While our study focused on adult mice (12–15 weeks old), Lu et al. characterized adolescent mice (7–8 weeks old). Additionally, Camp et al (2023) demonstrated that *Grin2a* ablation prolongs physiological maturation of hippocampal PV+ INs, suggesting synaptic differences in younger mice may not fully manifest.

### SST+ Interneurons and SCZ Pathology

Our data underscore the importance of SST+ INs in maintaining inhibitory tone and network stability, especially in upper cortical layers linked to SCZ risk (Duncan et al., 2024). These cells guide feedback processing and long-range gamma synchronization (Michalareas et al., 2016; Veit et al., 2017), which is critical for higher-order cognition. Thus, *Grin2a* ablation-induced SST+ dysfunction may help explain SCZ-related cognitive deficits. Although animal models cannot fully capture human SCZ complexity, they provide valuable mechanistic insights. Integrating animal findings with human studies will clarify the causal and translational significance of SST+ IN impairment in SCZ.

### Clinical and Translational Implications

Our use of *Grin2a* mutant mice as a model of NMDAR hypofunction offers a platform to study how disruptions in inhibitory circuits affect cortical network dynamics. While such models cannot capture the full spectrum of SCZ pathology, they allow us to investigate specific mechanisms—such as altered presynaptic calcium dynamics and enhanced gamma oscillations—that may underlie key disease phenotypes. Given the distinct effects of *Grin2a* ablation on PV+ and SST+ INs, therapeutic strategies may need to address both presynaptic and postsynaptic alterations. For PV+ INs, targeting presynaptic calcium dynamics with calcium channel modulators may normalize inhibitory control and gamma oscillations. For SST+ INs, postsynaptic receptor modulators could stabilize upper cortical layer networks.

### Study Limitations

While our study provides valuable insights, it is not without limitations. Phenotypic variability across *Grin2a* mouse models, including differences in electrophysiological properties and behavioral outcomes, underscores the complexity of studying NMDAR hypofunction. These inconsistencies may stem from temporal factors, such as developmental stages, or spatial constraints, including region- and cell-type-specific effects. Furthermore, while our findings suggest that altered presynaptic calcium dynamics drive asynchronous GABA release, direct measurements of presynaptic calcium levels were not performed. Future studies should address these gaps to validate the proposed mechanisms.

### Future Directions

Future research should investigate how disruptions in gamma oscillations during critical developmental periods contribute to neuropsychiatric disorders. For instance, studying the trajectory of gamma oscillatory power in *Grin2a* mutants across different developmental stages could elucidate how early alterations in inhibitory circuits shape long-term cortical and behavioral deficits. Such studies could also identify sensitive periods where interventions targeting GABAergic INs may most effectively mitigate disease progression. Understanding these trajectories may inform the design of therapies to restore normal oscillatory patterns during critical windows of brain development.

Exploring pharmacological agents targeting presynaptic calcium dynamics or modulating PV+ and SST+ IN activity could also offer new strategies for restoring inhibitory balance in SCZ. For example, presynaptic calcium channel modulators may address PV+ INs dysfunction, while postsynaptic receptor modulators could specifically target SST+ INs alterations. Additionally, non-pharmacological interventions, such as neuromodulation techniques, might complement pharmacological approaches by refining oscillatory dynamics and improving network stability. Future work should integrate developmental insights into therapeutic designs, tailoring interventions to align with the evolving state of inhibitory circuits and cortical networks.

## Conclusion

This study elucidates the critical role of *Grin2a-*encoded GluN2A-containing NMDARs in regulating the density and function of PV+ and SST+ INs within the mPFC. Using an animal model of NMDAR hypofunction relevant to SCZ, we demonstrate how *Grin2a* ablation disrupts inhibitory balance, enhances asynchronous GABA release, and amplifies gamma oscillations, particularly in upper cortical layers. These findings provide a mechanistic framework for understanding inhibitory circuit dysfunction in SCZ and highlight the need for translational research that bridges animal models with human studies.

## Acknowledgments

This work was supported by the Stanley Center for Psychiatric Research at the Broad Institute (KSJ). The authors report no biomedical financial interests or potential conflicts of interest. We thank Dr. Cagney Coomer for critical review of the manuscript.

## Author Contributions

K.S.J. conceptualized and designed the study. H.H. and S.E.M. conducted the study and performed analysis. K.S.J secured the funding. H.H. developed the methodology, and H.H. and S.E.M. carried out the experiments. H.H. constructed apparatuses. K.S.J. oversaw project administration and supervision. H.H. and K.S.J. wrote the original draft of the manuscript. All authors reviewed and edited the manuscript, contributed to interpreting the results, and provided critical feedback throughout the research process.

