## Supplemental_Files for "*Grin2a* Ablation Distinctly Reshapes PV+ and SST+ Interneuron circuits in the Medial Prefrontal Cortex to Amplify Gamma Oscillations"

**Supplementary Materials**

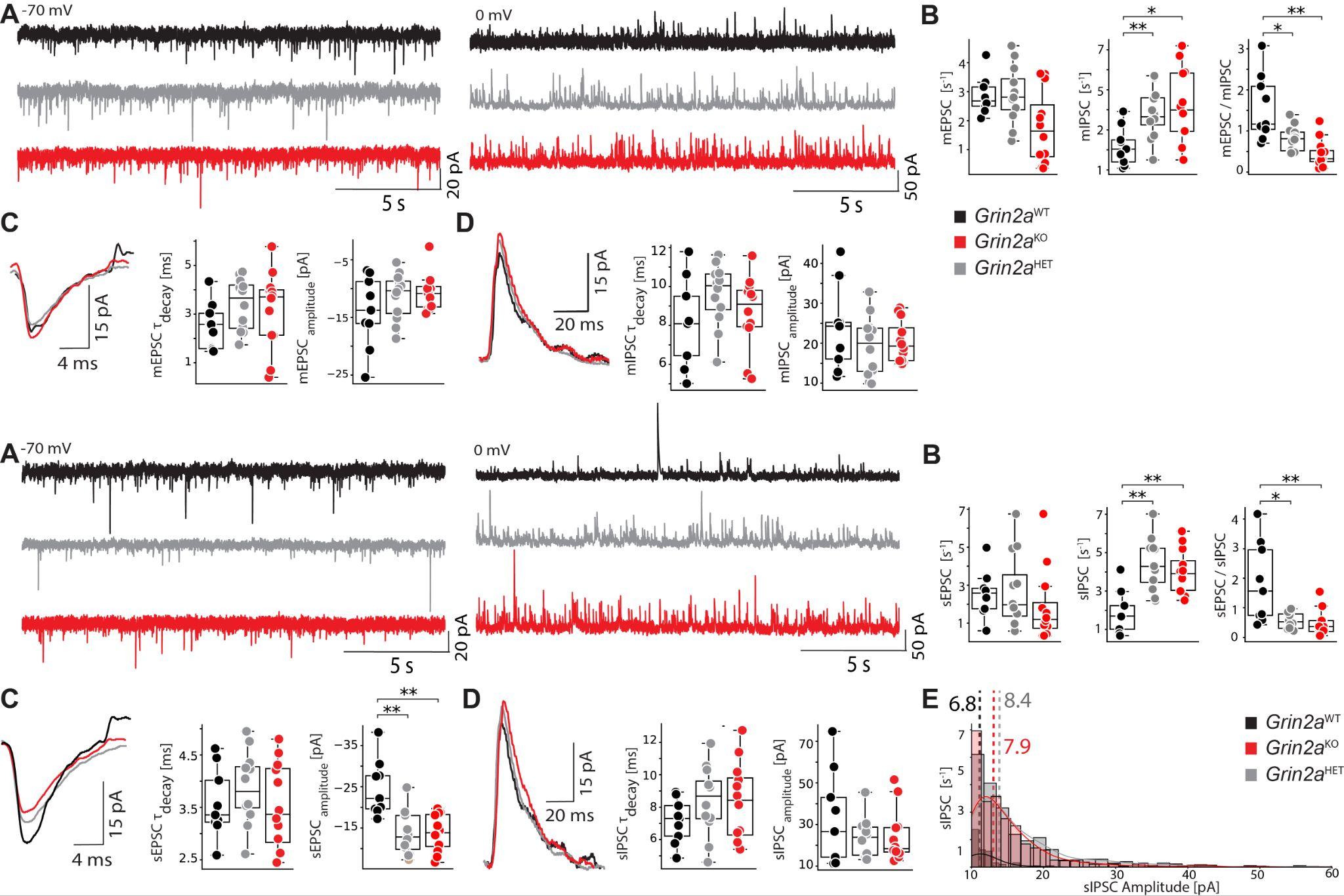

**Figure S1. Miniature Excitatory and Inhibitory Currents of *Grin2a* Variants in PrL**

(**A**) Representative traces showing miniature excitatory postsynaptic currents (mEPSCs) at −70 mV and miniature inhibitory postsynaptic currents (mIPSCs) at 0 mV in pyramidal neurons from different *Grin2a* genotypes. (**B**) Box plots illustrating the ratio of mIPSC to mEPSC frequencies across genotypes (WT: 0.8 ± 0.12; *Grin2a*^HET^, and *Grin2a*^HET^: 1.4 ± 0.15; *Grin2a*^HET^, and *Grin2a*^KO^: 3.7 ± 0.88). (**C**) Representative traces and quantification of mEPSC kinetics and amplitudes. Data represent mean ± SEM from 10–15 cells per genotype. Statistical significance was determined using the Kruskal-Wallis test: with p < 0.05 (*), p < 0.01 (**), p < 0.001 (***).

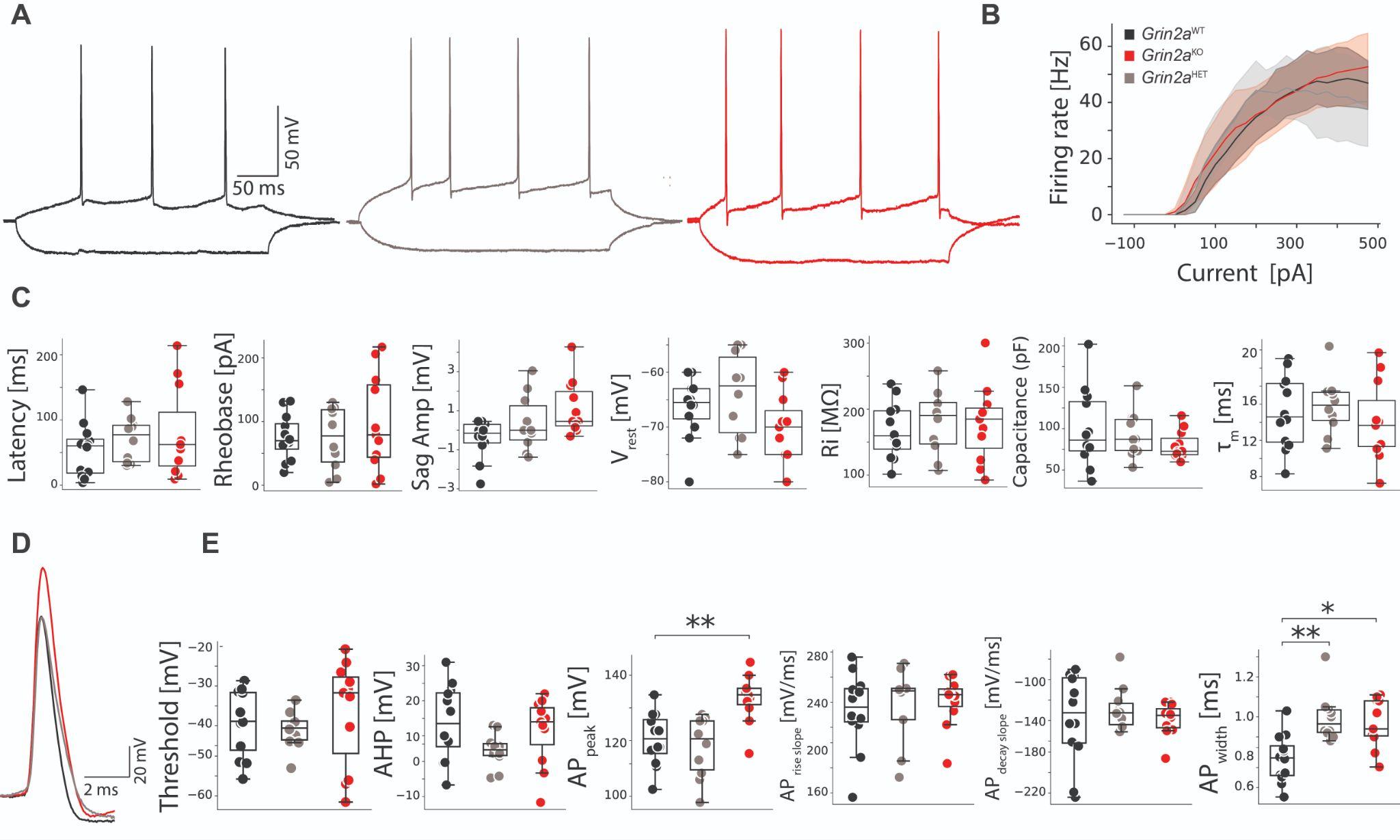

**Figure S2 Intrinsic Active Properties of L5 PrL Neurons of *Grin2a* Mutants in PrL**

**(A)** Representative traces showing evoked repetitive firing of pyramidal neurons in PrL cortical layer V of brain slices from wild-type and mutant mice. **(B)** Input-output (I-O) curves for AP firing of PrL pyramidal neurons of different mutants in response to current injections. I-O curves were generated by plotting the number of evoked APs by 300 ms current injections against current intensities over a range of −125 to 475 pA. **(C)** Quantification of different intrinsic passive properties of different mutant mice. **(D, E)** Representative AP waveform and kinetics quantification of different mutant mice.  Data represent mean ± SEM from 9-12 cells from 3 mice per genotype. Statistical significance was determined using the Kruskal-Wallis test, with p < 0.05 (*), and p < 0.01 (**).

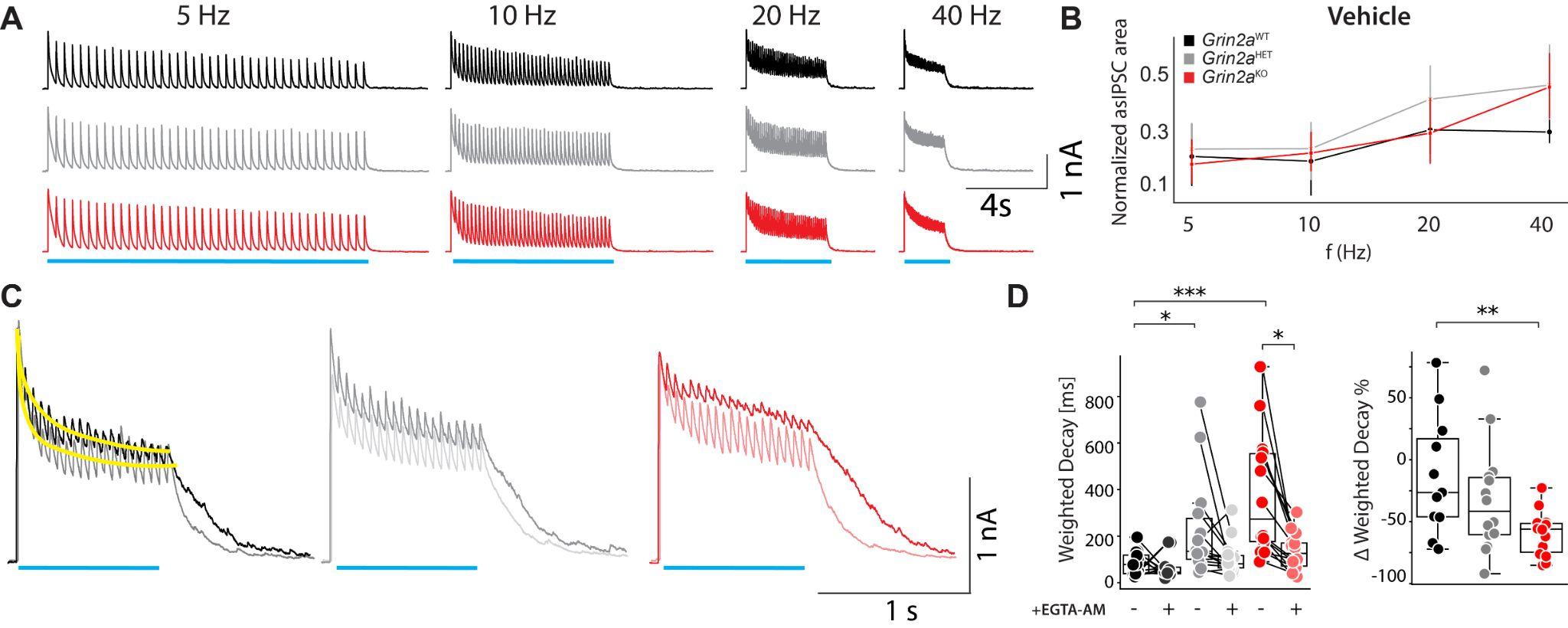

**Figure S3. Frequency-Dependent Asynchronous GABA Release from PV+ INs**

**(A)** Representative traces of tetanus-induced asynchronous IPSCs (asIPSCs) evoked by varying stimulation frequencies (5, 10, 20, 40 Hz) in WT (black), *Grin2a*^HET^ (gray), and *Grin2a*^KO^ (red) slices. Higher stimulation frequencies generally increase asynchronous release, and *Grin2a* mutants exhibit more pronounced asIPSCs. **(B)** Normalized post-tetanic asIPSC areas (averaged over 3 pre-tetanic IPSCs) as a function of stimulus frequency show that while WT slices display moderate increases at higher frequencies, *Grin2a*^HET^ and *Grin2a*^KO^ slices present significantly larger asIPSC areas at 20 Hz and 40 Hz. This indicates that GluN2A hypofunction heightens the frequency-dependent accumulation of residual presynaptic Ca²⁺ that drives asynchronous release. **(C)** Representative traces showing short-term depression during a high-frequency stimulus (40 Hz) in the absence (dark) or presence (light) of EGTA-AM. Traces were fit to a weighted tau (shown in yellow for WT). **(D)** Short-term depression emerges more slowly in *Grin2a* mutants. EGTA-AM accelerates short-term depression most dramatically in *Grin2a^KO^*, confirming that residual presynaptic Ca²⁺ underlies their excessive asynchronous release and a decrease short-term depression. Data represent mean ± SEM from 9–12 cells obtained from 3 mice per genotype. Statistical significance was determined using the Kruskal-Wallis test, with p < 0.05 (*) and p < 0.01 (**).

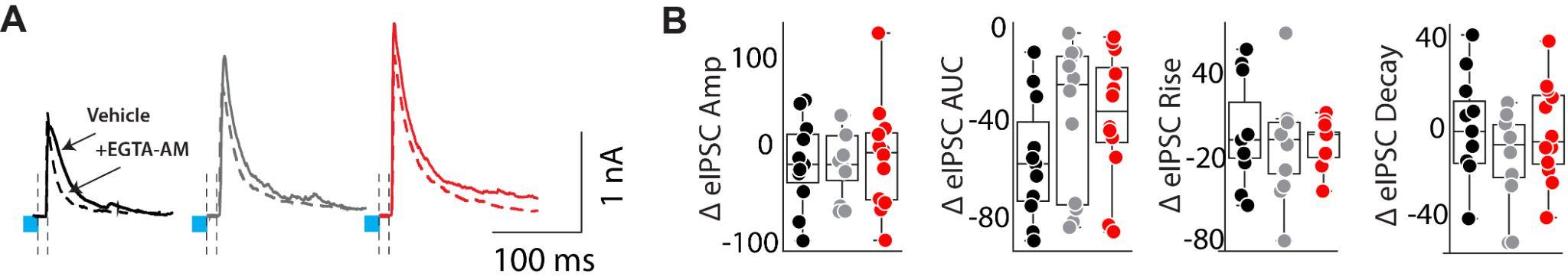

**Figure S4. Effect of EGTA-AM on SST-Driven Evoked IPSCs in Layer V Pyramidal Neurons**

(A) Representative averaged eIPSC traces from SST+ interneuron stimulation in WT (black), *Grin2a*^HET^ (gray), and *Grin2a*^KO^ (red) neurons before (solid lines) and after EGTA-AM application (dashed lines). EGTA-AM chelates presynaptic Ca²⁺, thereby assessing whether asynchronous GABA release contributes to SST-driven IPSC kinetics in each genotype.

(B) Summary of percent changes in eIPSC properties following EGTA-AM treatment, including peak amplitude, the area under the curve (AUC), rise time, and decay time. While WT neurons show minimal shifts, *Grin2a*^HET^ and *Grin2a*^KO^ neurons display more pronounced reductions in AUC and decay kinetics after EGTA-AM. This suggests that SST-mediated inhibition in mutants may have a component of Ca²⁺-dependent asynchronous release, albeit smaller than that observed in PV+ circuits (see Figures 3–4). Data represent mean ± SEM from 10–14 cells obtained from 3 mice per genotype**.**

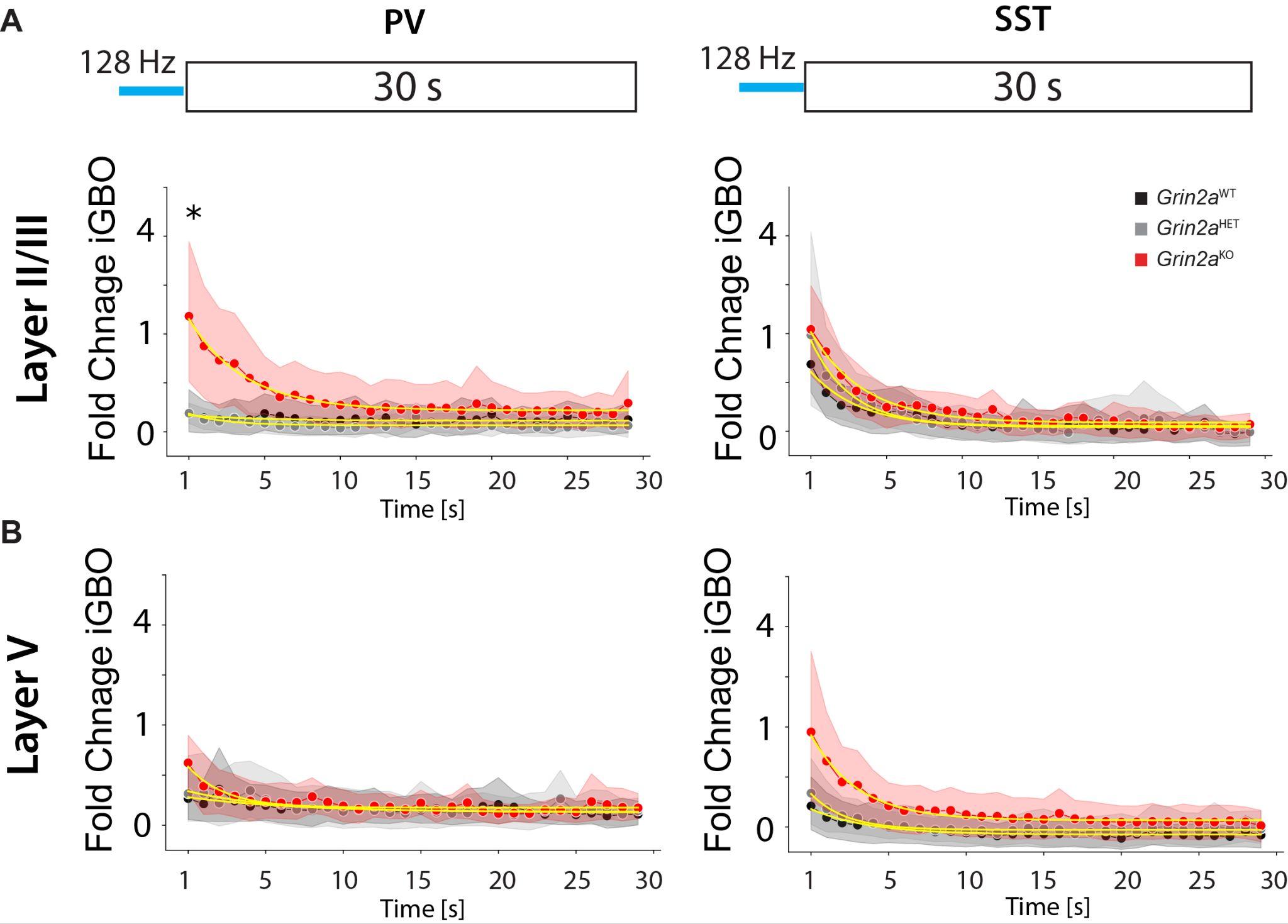

**Figure S5. Extended Dynamics of PV+ and SST+ iGBOs Following High-Frequency Stimulation**

(**A**) Time-course of iGBO (inhibitory gamma-band oscillation) power recorded from layer II/III of the mPFC up to 30 seconds after optogenetic stimulation of PV+ (left) or SST+ (right) interneurons at 128 Hz. WT (black), *Grin2a*^HET^ (gray), and *Grin2a*^KO^ (red) traces are shown. The weighted decay tau (yellow line) quantifies how quickly the induced oscillations return to baseline. For PV+ stimulation, *Grin2a*^KO^ slices exhibit a notably prolonged decay (WT: 2.06 ± 0.81 s, *Grin2a*^HET^: 2.97 ± 1.36 s, *Grin2a*^KO^: 5.86 ± 2.24 s), reflecting sustained gamma power. SST+ stimulation leads to less pronounced differences (WT: 4.84 ± 1.63 s, *Grin2a*^HET^: 4.22 ± 1.2 s, *Grin2a*^KO^: 5.42 ± 2.93 s).

(**B**) Similar recordings from layer V reveal that while both PV+ and SST+ stimulation can evoke iGBOs, the prolonged decay seen in *Grin2a*^KO^ mice is less marked compared to layer II/III. Weighted decay tau values for PV+ stimulation are WT: 3.49 ± 1.17 s, *Grin2a*^HET^: 3.85 ± 0.81 s, *Grin2a*^KO:^ 3.02 ± 0.7 s; and for SST+ stimulation, WT: 4.92 ± 1.59 s, *Grin2a*^HET:^ 3.9 ± 0.89 s, *Grin2a*^KO^: 4.41 ± 0.64 s.Data represent mean ± SEM from 8–9 mPFC slices obtained from 3 mice per genotype. Statistical significance was determined using the Kruskal-Wallis test, with p < 0.05 (*).

### **Table 1. Evoked LFP Amplitudes in Layer II/III of the mPFC Following PV+ Interneuron Stimulation at Multiple Frequencies** This table summarizes the mean local field potential (LFP) amplitudes (in mV) recorded from layer II/III of the medial prefrontal cortex (mPFC) slices obtained from WT, Grin2a^HET, and Grin2a^KO mice. Measurements were taken in response to optogenetic stimulation of PV+ interneurons at frequencies ranging from 1 to 128 Hz. Each value represents the mean ± SD, with N indicating the number of mice per group and n the number of slices analyzed.

| **Frequency (Hz)** | **WT (Mean ± SD, mV)**  ***(N = 3, n= 8-9)*** | ***Grin2a^HET^* (Mean ± SD, mV)**  ***(N = 3, n= 8-9)*** | ***Grin2a^KO^***  **(Mean ± SD, mV)**  ***(N = 3, n= 8-9)*** |
| --- | --- | --- | --- |
| 1 | -1.70 ± 0.31 | -1.58 ± 0.22 | -1.85 ± 0.23 |
| 2 | -1.34 ± 0.19 | -1.27 ± 0.14 | -1.36 ± 0.13 |
| 4 | -1.30 ± 0.15 | -1.29 ± 0.12 | -1.31 ± 0.10 |
| 8 | -1.25 ± 0.14 | -1.24 ± 0.08 | -1.29 ± 0.10 |
| 16 | -1.25 ± 0.13 | -1.36 ± 0.09 | -1.40 ± 0.13 |
| 32 | -1.26 ± 0.14 | -1.33 ± 0.07 | -1.39 ± 0.12 |
| 64 | -1.30 ± 0.14 | -1.30 ± 0.06 | -1.42 ± 0.12 |
| 128 | -1.31 ± 0.14 | -1.38 ± 0.05 | -1.42 ± 0.11 |

### **Table 2.** Evoked LFP Amplitudes (mV) in Layer V of the mPFC Following Optogenetic Stimulation of PV+ Interneurons. Values are presented as mean ± SD. *N* denotes the number of animals; *n* denotes the number of slices analyzed.

| **Frequency (Hz)** | **WT (Mean ± SD, mV)**  ***(N = 3, n= 8-9)*** | ***Grin2a^HET^* (Mean ± SD, mV)**  ***(N = 3, n= 8-9)*** | ***Grin2a^KO^* (Mean ± SD, mV)**  ***(N = 3, n= 8-9)*** |
| --- | --- | --- | --- |
| 1 | -1.94 ± 0.32 | -1.76 ± 0.24 | -1.86 ± 0.20 |
| 2 | -1.57 ± 0.18 | -1.43 ± 0.13 | -1.51 ± 0.14 |
| 4 | -1.52 ± 0.14 | -1.39 ± 0.13 | -1.46 ± 0.15 |
| 8 | -1.48 ± 0.15 | -1.36 ± 0.13 | -1.42 ± 0.13 |
| 16 | -1.48 ± 0.14 | -1.35 ± 0.10 | -1.45 ± 0.11 |
| 32 | -1.46 ± 0.14 | -1.36 ± 0.08 | -1.50 ± 0.14 |
| 64 | -1.46 ± 0.13 | -1.36 ± 0.08 | -1.50 ± 0.12 |
| 128 | -1.50 ± 0.13 | -1.42 ± 0.08 | -1.53 ± 0.12 |

**Table 3. Evoked LFP Amplitudes in Layer II/III of the mPFC Following SST+ Interneuron Stimulation at Multiple Frequencies**
This table presents mean ± SD values of local field potential (LFP) amplitudes (in mV) recorded from layer II/III of the mPFC in WT, *Grin2a*^HET^, and *Grin2a*^KO^ mice following optogenetic stimulation of SST+ interneurons at various frequencies (1–128 Hz). N indicates the number of animals and n is the number of slices tested per genotype. Significant differences from WT are denoted by *p < 0.05 and **p < 0.01 as determined by the Kruskal-Wallis test.

| **Frequency (Hz)** | **WT**  **(Mean ± SD, mV)  *(N = 3, n= 8-9)*** | ***Grin2a^HET^***  **(Mean ± SD, mV)**  ***(N = 3, n= 8-9)*** | ***Grin2a^KO^***  **(Mean ± SD, mV)**  ***(N = 3, n= 8-9)*** |
| --- | --- | --- | --- |
| 1 | -3.01 ± 0.25 | -5.17 ± 0.65 * | -3.51 ± 0.41 |
| 2 | -2.92 ± 0.24 | -4.93 ± 0.63 * | -3.66 ± 0.46 |
| 4 | -2.88 ± 0.23 | -4.82 ± 0.61 ** | -3.90 ± 0.45 |
| 8 | -2.88 ± 0.23 | -4.74 ± 0.65 * | -4.21 ± 0.54 |
| 16 | -2.99 ± 0.24 | -4.82 ± 0.71 | -4.30 ± 0.59 |
| 32 | -3.13 ± 0.27 | -4.87 ± 0.67 | -4.29 ± 0.59 |
| 64 | -3.07 ± 0.26 | -4.64 ± 0.67 | -3.86 ± 0.42 |
| 128 | -3.08 ± 0.26 | -4.63 ± 0.65 | -3.75 ± 0.42 |

**Table 4. Evoked LFP Amplitudes in Layer V of the mPFC Following SST+ Interneuron Stimulation at Multiple Frequencies**

This table lists the mean ± SD values of local field potential (LFP) amplitudes (mV) recorded from layer V of the mPFC in WT, Grin2a^HET, and Grin2a^KO mice after optogenetic stimulation of SST+ interneurons at frequencies ranging from 1 to 128 Hz. N indicates the number of animals per genotype, and n represents the number of slices analyzed. Statistical significance, determined by the Kruskal-Wallis test, is marked relative to WT values (*p < 0.05, **p < 0.01).

| **Frequency (Hz)** | **WT (Mean ± SD, mV)**  ***(N = 3, n= 8-9)*** | ***Grin2a^HET^* (Mean ± SD, mV)**  ***(N = 3, n= 8-9)*** | ***Grin2a^KO^* (Mean ± SD, mV)**  ***(N = 3, n= 8-9)*** |
| --- | --- | --- | --- |
| 1 | -2.99 ± 0.28 | -5.07 ± 0.57 * | -3.73 ± 0.47 |
| 2 | -2.68 ± 0.25 | -4.45 ± 0.43 ** | -3.37 ± 0.43 |
| 4 | -2.55 ± 0.24 | -4.37 ± 0.38 ** | -3.46 ± 0.42 |
| 8 | -2.49 ± 0.24 | -4.23 ± 0.34 ** | -3.56 ± 0.45 |
| 16 | -2.48 ± 0.24 | -4.18 ± 0.41 ** | -3.64 ± 0.48 |
| 32 | -2.66 ± 0.24 | -4.03 ± 0.38 * | -3.42 ± 0.40 |
| 64 | -2.64 ± 0.20 | -3.91 ± 0.36 * | -3.33 ± 0.37 |
| 128 | -2.62 ± 0.18 | -4.09 ± 0.39 * | -3.27 ± 0.38 |
